# On Improving an Already Competitive Segmentation Algorithm for the Cell Tracking Challenge - Lessons Learned

**DOI:** 10.1101/2021.06.26.450019

**Authors:** Tim Scherr, Katharina Löffler, Oliver Neumann, Ralf Mikut

## Abstract

The virtually error-free segmentation and tracking of densely packed cells and cell nuclei is still a challenging task. Especially in low-resolution and low signal-to-noise-ratio microscopy images erroneously merged and missing cells are common segmentation errors making the subsequent cell tracking even more difficult. In 2020, we successfully participated as team KIT-Sch-GE (1) in the 5th edition of the ISBI Cell Tracking Challenge. With our deep learning-based distance map regression segmentation and our graph-based cell tracking, we achieved multiple top 3 rankings on the diverse data sets. In this manuscript, we show how our approach can be further improved by using another optimizer and by fine-tuning training data augmentation parameters, learning rate schedules, and the training data representation. The fine-tuned segmentation in combination with an improved tracking enabled to further improve our performance in the 6th edition of the Cell Tracking Challenge 2021 as team KIT-Sch-GE (2).

## 1 Introduction

Cell segmentation is a fundamental step in many developmental biology studies. It enables the study of cell behavior in developmental processes such as tissue development and organ formation and therefore gives insight into crucial biological processes at cell level [1, 2, 3]. However, due to the large amount of data generated with modern microscopy techniques, automated, easy-to-use, and accurate segmentation methods are needed. In recent years, deep learning-based segmentation methods have proven to perform well for many imaging techniques and cell morphologies [4, 5, 6, 7]. An advantage of deep learning methods is that the required expert knowledge is shifted towards the training process, enabling to build easy-to-use-tools for biologists, when the trained models generalize well.

Deep learning-based instance segmentation methods can be roughly categorized into: (i) semantic methods with multiclass predictions, e.g., background, cell and cell boundary, to get instance information [8, 9], (ii) methods predicting distance or gradient flow maps [6, 10], (iii) proposal-based methods like region proposals [11] or polygon proposals [12], (iv) instance embedding methods predicting similar embeddings for pixels belonging to the same object and dissimilar embeddings otherwise [13], and (v) a combination of aforementioned methods, e.g., predicting gradient flow maps and cell regions, i.e., the two classes background and cell [7]. Often, adapted loss functions are used, e.g., to improve the class separation in multiclass prediction methods [14].

The large number of deep learning-based methods, which have all proven to work well at least for a specific scenario, makes it difficult to find the best method in combination with a good parametrization for the data to evaluate. However, we think that it is often easier to improve the training data set and training process of an already implemented and working method. In 2020, we successfully participated as team KIT-Sch-GE (1) in the ISBI Cell Tracking Challenge^1^ with a distance map regression method [6].

We achieved multiple top 3 rankings on diverse data sets showing different cell types and imaging techniques. In this manuscript, we show that simple training and method refinements can lead to a significant improvement in segmentation quality on a rather small training data set. With the improved segmentation, newly created training data sets and a new tracking [15], we were able to further improve our results in the Cell Tracking Challenge as team KIT-Sch-GE (2).

## 2 Materials and Method

For the segmentation, a U-Net with two decoder paths is trained to predict cell distance maps and neighbor distance maps [6]. The cell distance represents the normalized Euclidean distance of each pixel of a cell to the nearest pixel not belonging to this cell (Figure 1c). The neighbor distance represents an inverse normalized distance of a cell’s pixel to the closest pixel of another cell (Figure 1d). This information about neighbors should further prevent undersegmentation errors. The cell and the neighbor distance are normalized to the range [0, 1] for each cell. In the post-processing, both predictions are smoothed slightly, the threshold *th*_mask_ is applied to adjust the cell size and the threshold seed *th*_seed_ is applied to get seeds for a watershed segmentation. Our segmentation method is described in detail and compared with other cell segmentation approaches in [6]. The code of our fine-tuned segmentation is available at: https://git.scc.kit.edu/KIT-Sch-GE.

**Figure 1.**
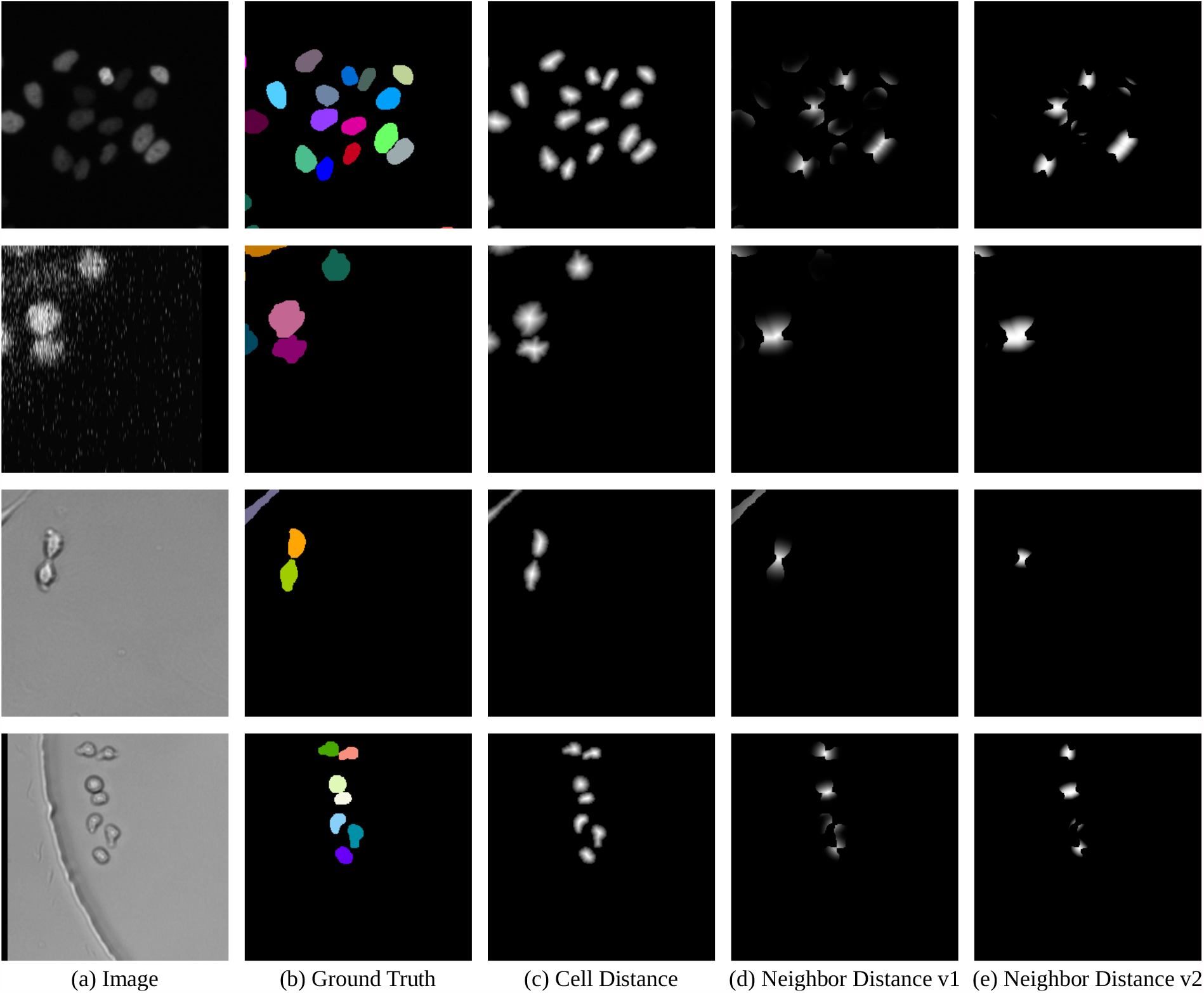
Training Data Set Examples. Shown are raw images (a), ground truths (b), cell distance maps (c), the neighbor distance maps from [6] (d), and the fine-tuned neighbor distance maps (e) of the four in the training data set included cell types Fluo-N2DL-HeLa (1st row), Fluo-N3DH-CE (2nd row), BF-C2DL-MuSC (3rd row) and BF-C2DL-HSC (4th row). Touching regions are more distinct in the fine-tuned neighbor distance maps than in the original version. Moreover, the fine-tuned neighbor distance maps show less normalization artifacts. Images and ground truths are crops from Cell Tracking Challenge data [16, 17].

### 2.1 Neighbor Distance v2

During the creation of the neighbor distance v1 proposed in [6], a scaling step is applied (Figure 3h in [6]). In the fine-tuned version, referred to as neighbor distance v2, a custom scaling function is applied, which is steeper for small values and less steep for high values resulting in more distinct touching regions. Instead of the grayscale closing (Figure 3g in [6]), a bottom-hat-transform closing is applied to close small gaps. Figure 1e shows the fine-tuned neighbor distance v2 for some cell types of the Cell Tracking Challenge [16, 17]. Further, in the normalization step, the minor axis length of the cell is incorporated reducing some artifacts for elongated cells (see Figure 1d&e 3rd row).

**Figure 2.**
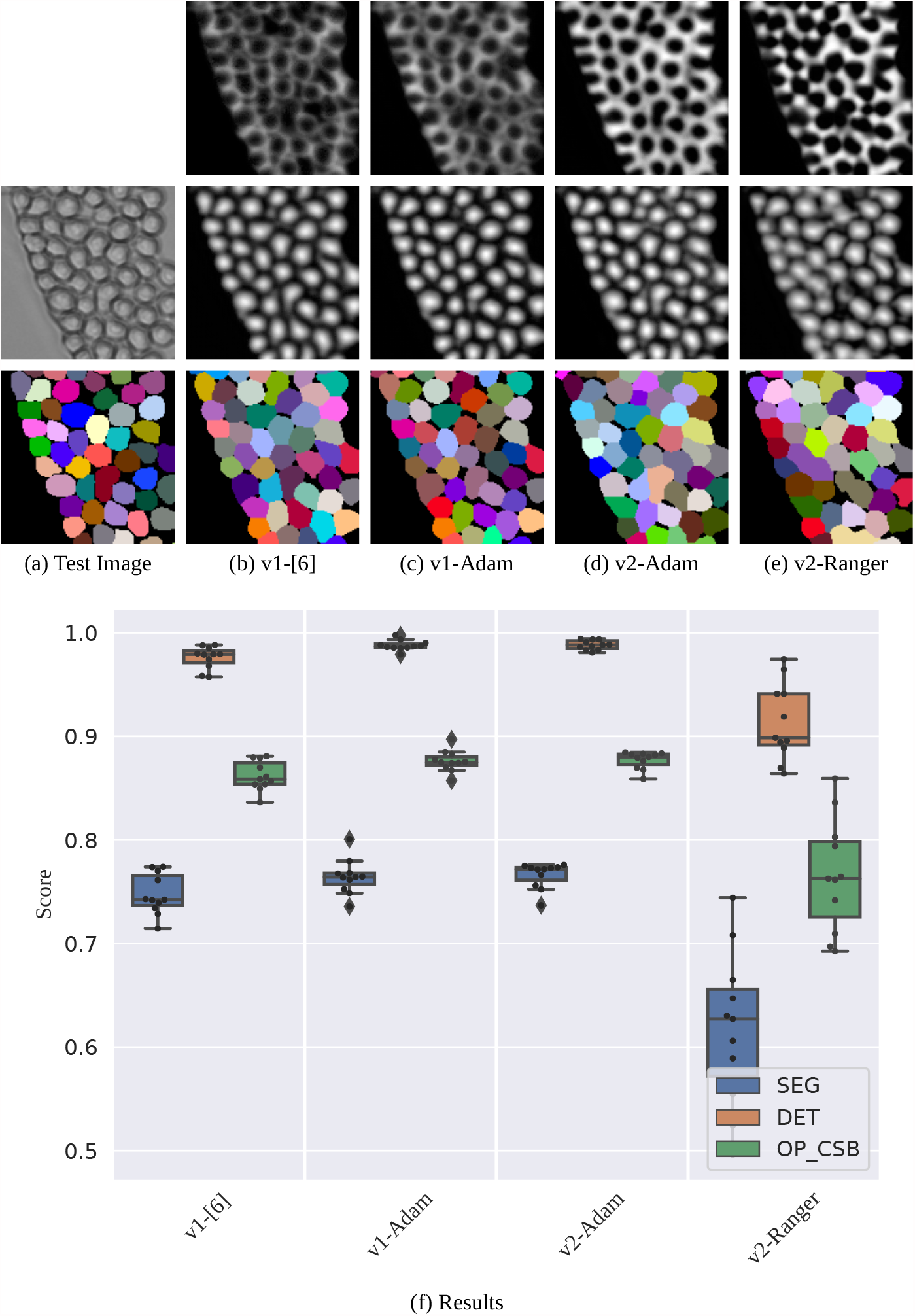
Segmentation Results on the BF-C2DL-HSC Test Set. Shown are raw images and segmentations of a 140 px×140 px test image crop (a-e, best OP_CSB_ models). The plot on the bottom shows the evaluation on the test set with 11 initializations per method (f).

**Figure 3.**
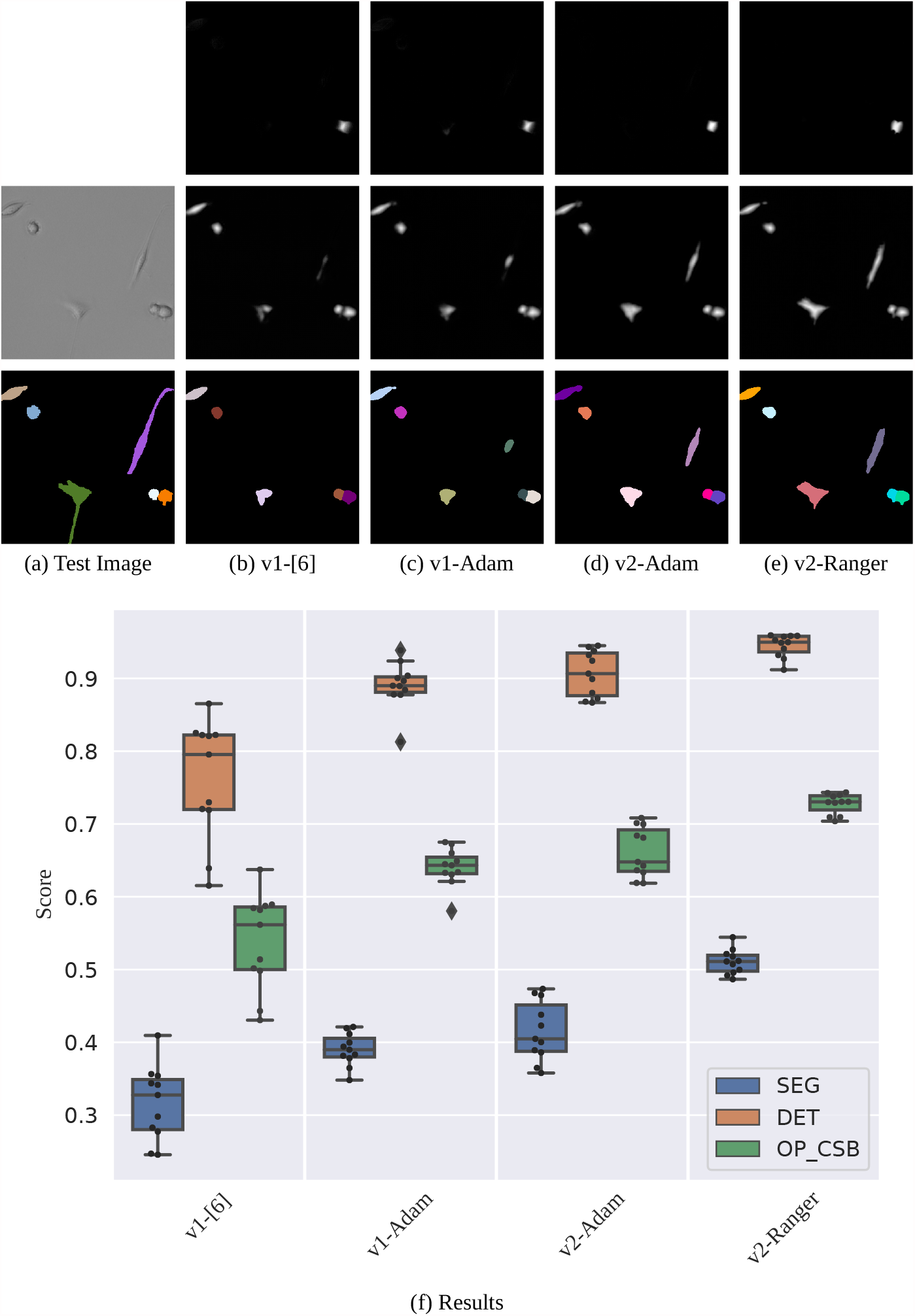
Segmentation Results on the BF-C2DL-MuSC Test Set. Shown are raw images and segmentations of a 140 px×140 px test image crop (a-e, best OP_CSB_ models). The plot on the bottom shows the evaluation on the test set with 11 initializations per method (f).

### 2.2 Training Process Adjustments

In [6] and for our Cell Tracking Challenge contribution in 2020, models were trained for a maximum number of epochs 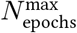 with the Adam optimizer [18] and the start learning rate *lr*_start_. The loss function was PyTorch’s SmoothL1Loss and a batch size of 8 has been used. After a patience *N*_patience_ of subsequent epochs without validation loss improvement, the learning rate has been multiplied with 0.25 till the minimum learning rate *lr*_min_ was reached. An early stopping criterion, which stops the training process after *N*_stop_ subsequent epochs without validation loss improvement, has been applied. For inference, the best validation loss model is used. Furthermore, training data augmentation has been applied.

#### 2.2.1 Training Settings

Table 1 shows the fine-tuned settings for the training data set used in [6] (see also Section 2.4). The learning rates have been adapted by analyzing a short run of only a few epochs with increasing learning rate which is sufficient to estimate boundary learning rates [19]. Furthermore, we increased the patience *N*_patience_ which required increased parameters 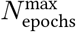 and *N*_stop_ as well.

**Table 1.** Training Settings. We found that for Adam 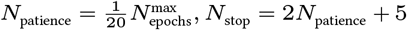 works well, and for Ranger 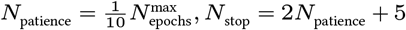. In our experience, A larger patience does only very rarely improve the validation loss but increases the training time significantly.

| Parameter | Training in [6] | Fine-Tuned Training |  |  |
| --- | --- | --- | --- | --- |
|  |  | Adam | Ranger | Ranger (2nd run) |
| $N_{\text{epochs}}^{\text{max}}$ | 200 | 320 | 320 | 32 |
| $N_{\text{patience}}$ | 12 | 16 | 32 | - |
| $N_{\text{stop}}$ | 28 | 37 | 69 | - |
| $lr_{\text{start}}$ | $8 \times 10^{-4}$ | $8 \times 10^{-4}$ | $6 \times 10^{-3}$ | $5.4 \times 10^{-4}$ |
| $lr_{\text{min}}$ | $6 \times 10^{-5}$ | $3 \times 10^{-6}$ | $4.5 \times 10^{-4}$ | $3 \times 10^{-5}$ |

#### 2.2.2 Ranger Optimizer

Ranger is an optimizer which combines Rectified Adam [20], Lookahead [21], and gradient centralization [22] in one optimizer^2^. Rectified Adam provides an automated warm-up to ensure a solid start to training [20]. Lookahead iteratively updates two sets of weights [21] and chooses a search direction by looking ahead at the sequence of ”fast weights” generated by another optimizer. We use Ranger with the training settings provided in Table 1. After an initial training run, the best validation loss model is further trained in a second run with cosine annealing [23].

#### 2.2.3 Training Data Augmentation

The training data set in [6] has been augmented using the following augmentation sequence with augmentation probabilities given in brackets:

- flipping (75%): up-down flip, left-right flip or 90° rotation.
- scaling (30%): scale factor *s* ∈ [0.85, 1.15].
- rotation (30%): angle *α* ∈ [−180°, 180°].
- contrast (30%): CLAHE (openCV) or mean & gamma changes.
- blur (30%): Gaussian blur with random *σ* ∈ [0, 5].
- noise (30%): additive Gaussian noise with *σ* ∈ [1, 7] × max *I*_train_.

We adapted the sequence of augmentations and fine-tuned the parameters manually:

- flipping (100%): original image, left-right flip, up-down flip, 90° rotation, 180° rotation, 270° rotation, left-right flip + 90° rotation or up-down flip + 90° rotation.
- contrast (50%): CLAHE (skimage), percentile normalization or mean & gamma changes.
- scaling (25%): scale factor *s* ∈ [0.80, 1.20].
- rotation (25%): angle *α* ∈ [−45°, 45°].
- blur (30%): Gaussian blur with random *σ* ∈ [1, 2.75].
- noise (30%): additive Gaussian noise with *σ* ∈ [1, 6] × max *I*_train_.

Now the augmentations flipping and rotation can not undo each other, e.g., a 90° rotation cannot be reversed by the rotation augmentation. Most changes are small and the largest change is the maximum amount of blur that can be applied which could result in complete loss of inner cell structures. However, it is not clear if this improves the generalization ability or interferes the training process even though being applied very rarely.

### 2.3 Post-Processing

The use of the neighbor distance v2 and the improved training process enables the reduction of smoothing in the post-processing and leads to the same parameter value applicable for all cell types, whereas KIT-Sch-GE (1) used a per cell type adapted smoothing. To further improve the scaling, we apply the tangent function to the squared neighbor distances additionally. Instead of removing only very small seeds below a fixed threshold area, seeds smaller than 10% of the mean seed size in a frame are removed. If no cell is detected in a frame, the seed threshold is decreased iteratively till a seed is found or a lower bound is reached.

### 2.4 Training Data Set

We use the same training data set as in [6]. The data set consists of 268 crops of size 256 px×256 px and is created from the Cell Tracking Challenge data sets Fluo-N2DL-HeLa (Figure 1 1st row, 123 crops, 1404 cells), Fluo-N3DH-CE (Figure 1 2nd row, 18 crops, 69 cells), BF-C2DL-MuSC (Figure 1 3rd row, 100 crops, 174 cells) and BF-C2DL-HSC (Figure 1 4th row, 27 crops, 194 cells). The unequal distribution of the training images, i.e., of the cell types and of the cell density within the crops, makes this training data set challenging.

## 3 Results

All experiments were performed on a system with two NVIDIA TITAN RTX GPUs and PyTorch has been used as deep learning framework.

### 3.1 Data Set and Evaluation Criteria

Each cell type included in the training data set is evaluated separately on a by the Cell Tracking Challenge provided second set not used for the model training. A difficulty is that in the evaluation the segmentation of touching cells is very important since the cell number and density grow over time. In contrast, the used training set contains many samples from early time points.

The performance measures DET and SEG from the Cell Tracking Challenge are used. The DET metric evaluates object level segmentation errors, i.e., the adding of cells, the missing of cells and the merging of cells [24]. The SEG metric evaluates pixel level errors, i.e., how well ground truth and predicted masks do match. The overall performance for the Cell Segmentation Benchmark of the Cell Tracking Challenge is calculated as follows:

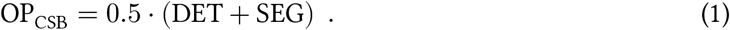

### 3.2 Compared Methods

We compare the methods

i. v1-[6]: cell distances & neighbor distances v1 (taken from [6], old training process),
ii. v1-Adam: cell distances & neighbor distances v1, Adam optimizer,
iii. v2-Adam: cell distances & neighbor distances v2, new post-processing, Adam optimizer,
iv. v2-Ranger: cell distances & neighbor distances v2, new post-processing, Ranger optimizer. For each method, eleven models are trained to evaluate the robustness.

### 3.3 Parameter Selection

For the methods (i) and (ii), we use the post-processing parameters from [6]: *th*_mask_ = 0.09, *th*_seed_ = 0.5. For the methods (iii) and (iv), we use the parameters: *th*_mask_ = 0.07, *th*_seed_ = 0.45. The mask threshold is lowered due to the smaller amount of smoothing in the new post-processing.

### 3.4 Influence of the Early Stopping Criterion

First, we evaluate the influence of the maximum amount of training epochs 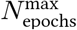 on the methods trained with the Adam optimizer to validate that improvements are not just due to an unfinished training process in [6]. For method (i) v1-[6], five of the eleven trained models stopped early, with the smallest amount of trained epochs of 130. For method (ii) v1-Adam all models stopped early with the smallest training time of 204 epochs. All trained models of method (iii) v2-Adam stopped early, with the smallest amount of trained epochs of 218. As almost half of the models of method (i) stopped early, we reason that increasing the maximum amount of epochs would not have resulted in relevant improvements to the results reported in [6]. Furthermore, we conclude that the longer training times are due to the adjusted learning rate schedule and training data augmentation.

### 3.5 BF-C2DL-HSC

The segmentation results on the BF-C2DL-HSC test set in Figure 2 show an improved DET score when applying the new training process (v1-Adam vs. v1-[6]). An improved DET score means that less false positives, false negatives and merges occur. This better object detection also has an influence on the pixel level evaluation SEG score, which is also slightly improved. The fine-tuned method shows better raw neighbor distance predictions when trained with the Adam optimizer, however, the scores do not further improve (v2-Adam vs. v1-Adam). Method (iv) trained using the Ranger optimizer does not perform competitively on this data set.

### 3.6 BF-C2DL-MuSC

On the data set BF-C2DL-MuSC, the new training process shows a large improvement in all scores (Figure 3). This is mainly due the better detection of the elongated cells. However, the fine-tuned version v2 even further improves the segmentation of the elongated cells, especially when trained with the Ranger optimizer. This is due to the better prediction of the cell distances. As the methods v1-Adam and v2-Adam do only differ in the neighbor distances and in the post-processing, the neighbor distances need to have an important influence on the training of the cell distances.

### 3.7 Fluo-N2DL-HeLa

Figure 4 shows that all methods perform quite similar on the data set Fluo-N2DL-HeLa. However, the new training process is more stable resulting in a narrower distribution of the single initializations. Again v2-Ranger yields the best results. In addition, the v2-Ranger models have the smallest variance.

**Figure 4.**
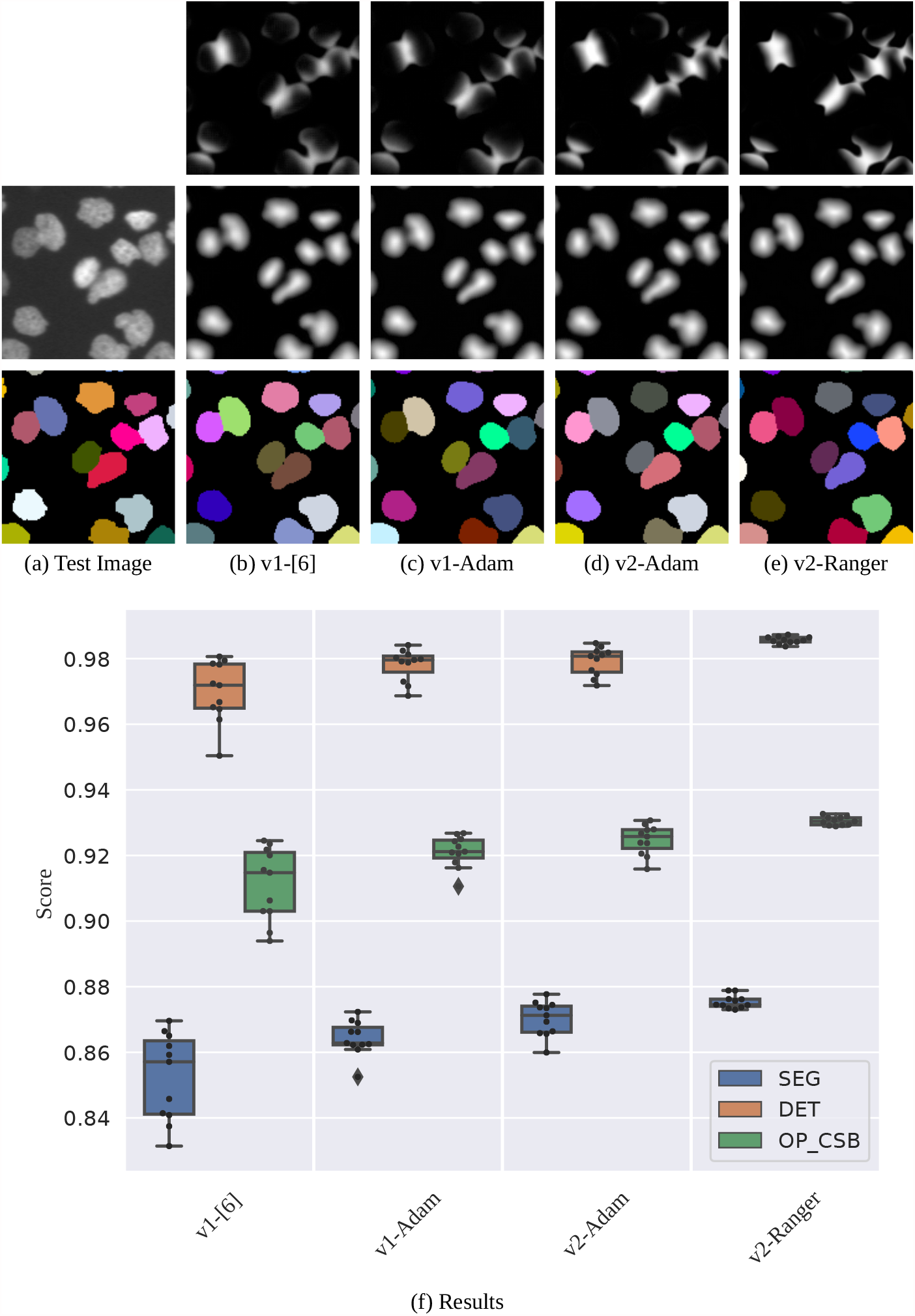
Segmentation Results on the Fluo-N2DL-HeLa Test Set. Shown are raw images and segmentations of a 140 px×140 px test image crop (a-e, best OP_CSB_ models). The plot on the bottom shows the evaluation on the test set with 11 initializations per method (f).

The segmentation error analysis in Figure 5 shows that the fine-tuned version v2-Adam shows less merged cells than v1-Adam, i.e., less splitting operations need to be applied to reach a perfect segmentation. v2-models trained with the Ranger optimizer predict fewer false negatives and merges but tend to oversegment. As oversegmentation errors are easier to correct in the tracking than undersegmentation errors, they are penalized less in the DET metric. The raw neighbor distance predictions in Figure 4 show that the use of the neighbor distance v2 results in more distinct touching regions.

**Figure 5.**
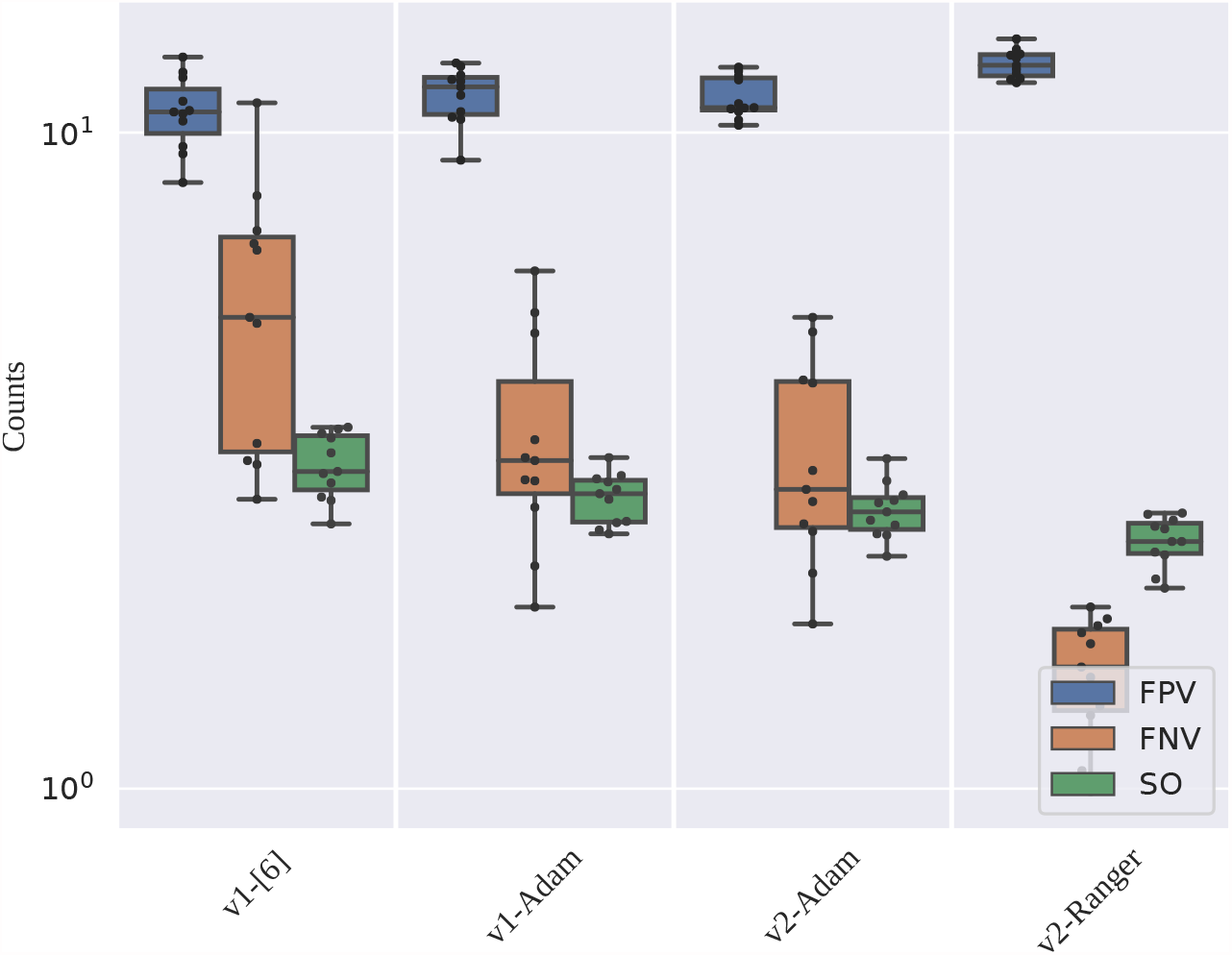
Segmentation Error Analysis for the Data Set Fluo-N2DL-HeLa. The plot shows the error counts per frame on the test set, FPV: false positive vertices, FNV: false negative vertices, SO: needed splitting operations. The data set shows on average 276 cells per frame and FPs and SOs can also occur due to too early and too late detected cell mitosis. The FPs also depend on a needed field of interest correction.

### 3.8 Fluo-N3DH-CE

On the 3D data set Fluo-N3DH-CE, the old training process used for v1-[6] shows the best results (Figure 6). However, since the predictions are made slice-wise and only the post-processing is 3D, a better training process and better 2D predictions do not necessarily mean a better 3D cell detection. The biggest problem on this data set are merged objects in axial direction. In our Cell Tracking Challenge contributions, we therefore used an iterative splitting scheme (see also [25]).

**Figure 6.**
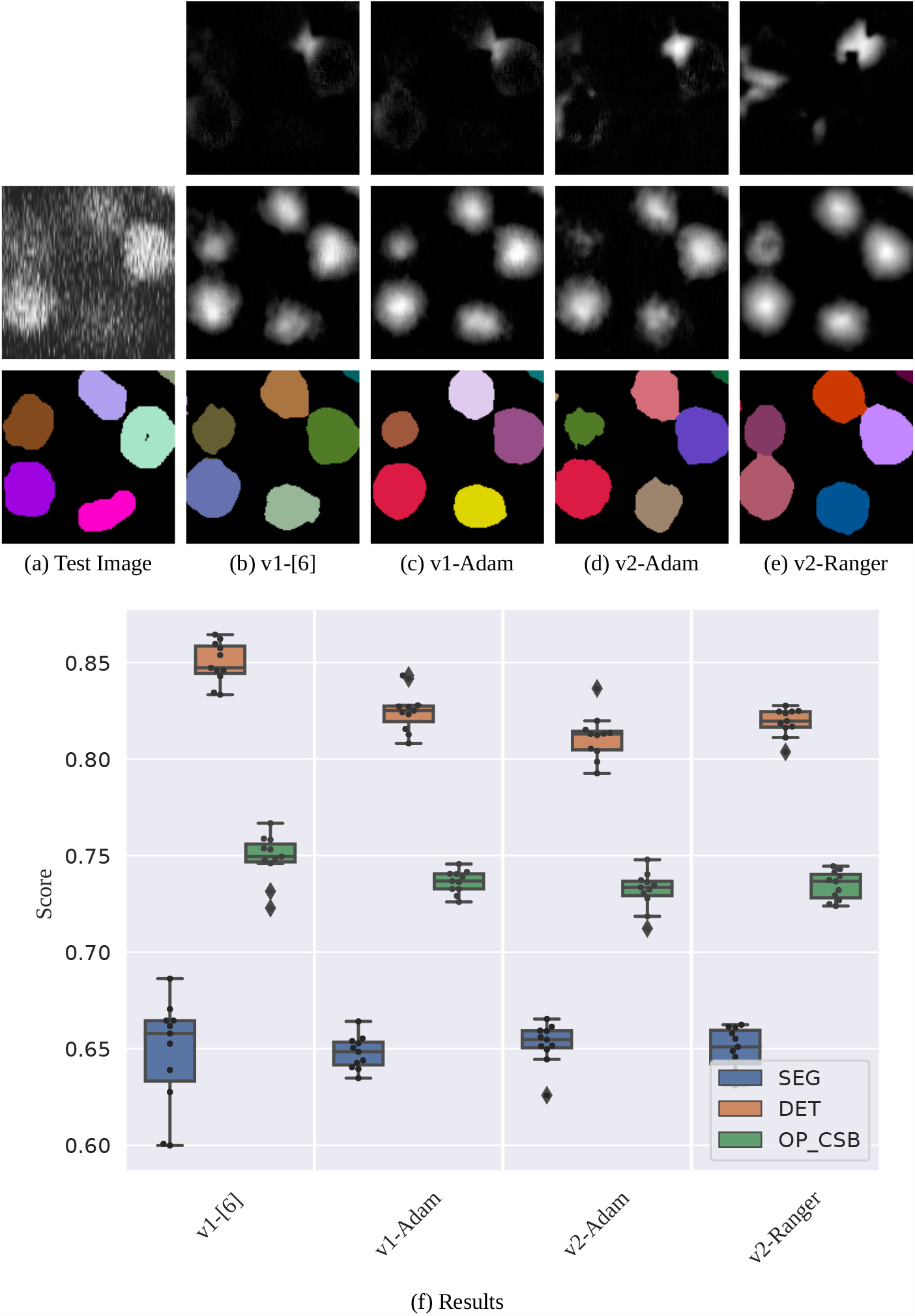
Segmentation Results on the Fluo-N3DH-CE Test Set. Shown are raw images and segmentations of a 140 px×140 px test image crop (a-e, best OP_CSB_ models). The plot on the bottom shows the evaluation on the test set with 11 initializations per method (f).

### 3.9 Cell Tracking Challenge

Table 2 shows the segmentation results of our submission for the 6th edition of the Cell Tracking Challenge 2021. Especially for 2D data sets with many touching cells, i.e., BF-C2DL-HSC (bright field), Fluo-N2DL HeLa (fluorescence) and PhC-C2DL-PSC (phase contrast), our segmentation method achieves good results. For more details, see [25], e.g., how the best models were selected, and [15] for our cell tracking.

**Table 2.**
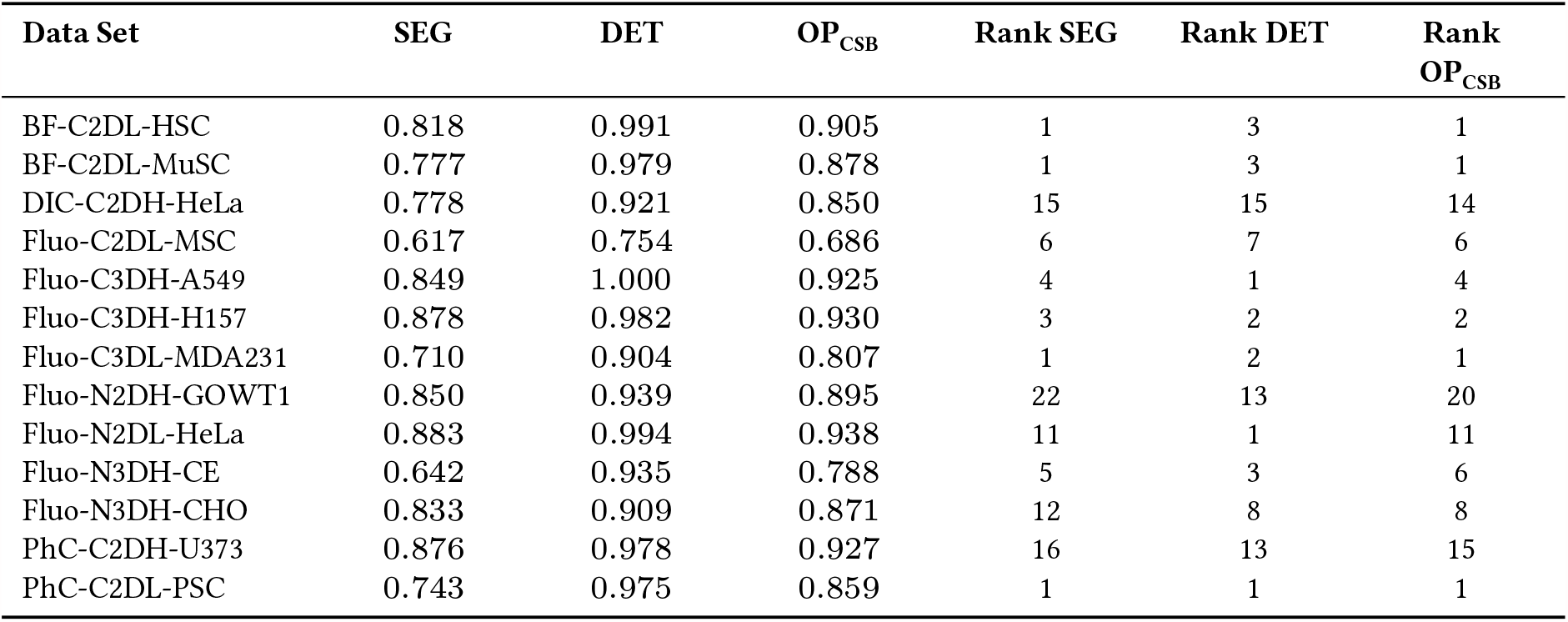
Cell Segmentation Benchmark Results (6th Cell Tracking Challenge Edition). The latest leaderboard is available on the Cell Tracking Challenge website. State of the results: April 13th 2021.

## 4 Discussion

Fine-tuning the training process enabled to further improve the segmentation performance on three out of the four in our training data set included cell types. For the 3D data set, a better training process and better 2D predictions do not necessarily mean a better 3D cell detection which could explain the drop in performance on that data set. The fine-tuned neighbor distances v2 show better raw predictions and also improved the median OP_CSB_ on the data sets Fluo-N2DL-HeLa and BF-C2DL-HSC. However, we observed the largest performance boost in the improved segmentation of the elongated cells in the data set BF-C2DL-MuSC. This is surprising since the cell distances were not fine-tuned. Maybe the neighbor distances v2 propagate more and better information in the shared encoder path during training.

Using the Ranger optimizer improves the segmentation of elongated cells further and reduces missing and merged cells on the Fluo-N2DL-HeLa data set. However, the drop in performance for the data set BF-C2DL-HSC when using the Ranger optimizer is interesting since the cells look quite similar to the BF-C2DL-MuSC cells in the not elongated state. This could be due to the fact that only 27 crops of the data set BF-C2DL-HSC are included in the training data set but 123 Fluo-N2DL-HeLa crops and 100 BF-C2DL-MuSC crops. Thus, Adam and Ranger seem to find different minima or saddle points during the optimization process.

The Cell Tracking Challenge results show that our fine-tuned method is competitive, especially for 2D data sets with many touching cells. Further, the results show that our method works well for bright field, phase contrast, and fluorescence images.

## 5 Conclusion and Outlook

In this manuscript, we have shown that simple fine-tuning of training process settings and augmentations can improve the segmentation performance significantly. In addition, minor changes to the training data representations, i.e., of the neighbor distances, also helped to improve the segmentation performance, especially for some specific cell shapes. Our results imply that the optimizers Adam and Ranger behave quite different in our setting and therefore complement each other well. We suggest to train models with Adam and with Ranger and not to stick to one optimizer.

Moreover, our results also show that selecting a well-parametrized augmentation sequence and learning rate schedules can be crucial to improve the performance. Automating the training process for instance by using tools like FasterAutoAugment [26] could lead to further performance boosts or reduced engineering times in the future.

## Acknowledgments

We would like to thank Andreas Bartschat for proofreading.

## Author Contributions

**Conceptualization** — TS, RM

**Methodology** — TS, RM

**Software** — TS, KL

**Writing - original draft** — TS

**Writing - review & editing** — TS, KL, ON, RM

**Funding acquisition** — RM

**Project administration** — RM

**Cell Tracking Challenge participation** — TS, KL

## Funding

We are grateful for the funding by the Helmholtz Association in the programs Natural, Artificial and Cognitive Information Processing (TS, RM) and the Helmholtz Information & Data Science School for Health (KL, RM). The funders had no role in study design, data collection and analysis, decision to publish, or preparation of the manuscript.

## Footnotes

1 http://celltrackingchallenge.net/

2 https://github.com/lessw2020/Ranger-Deep-Learning-Optimizer

## Notes

### Competing Interest Statement

The authors have declared no competing interest.

## References

[1] K. Khairy and P. J. Keller. “Reconstructing Embryonic Development”. In: Genesis 49 (2011), pp. 488–513. doi: 10.1002/dvg.20698.

[2] E. Meijering. “Cell Segmentation: 50 Years Down the Road [Life Sciences]”. In: IEEE Signal Process Mag 29.5 (2012), pp. 140–145. doi: 10.1109/MSP.2012.2204190.

[3] A. Y. Kobitski et al. “An Ensemble-Averaged, Cell Density-Based Digital Model of Zebrafish Embryo Development Derived from Light-Sheet Microscopy Data with Single-Cell Resolution”. In: Sci Rep 5.8601 (2015), pp. 1–10. doi: 10.1038/srep08601.

[4] J. C. Caicedo et al. “Nucleus Segmentation Across Imaging Experiments: The 2018 Data Science Bowl”. In: Nat Methods 16 (2019), pp. 1247–1253. doi: 10.1038/s41592-019-0612-7.

[5] R. Hollandi et al. “nucleAIzer: A Parameter-free Deep Learning Framework for Nucleus Segmentation Using Image Style Transfer”. In: Cell Syst 10 (2020), pp. 453–458. doi: 10.1016/j.cels.2020.04.003.

[6] T. Scherr et al. “Cell Segmentation and Tracking using CNN-Based Distance Predictions and a Graph-Based Matching Strategy”. In: PLoS One 15.12 (2020), pp. 1–22. doi: 10.1371/journal.pone.0243219.

[7] C. Stringer et al. “Cellpose: A Generalist Algorithm for Cellular Segmentation”. In: Nat Methods 18 (2021), pp. 100–106. doi: 10.1038/s41592-020-01018-x.

[8] O. Ronneberger, P. Fischer, and T. Brox. “U-Net: Convolutional Networks for Biomedical Image Segmentation”. In: Med Image Comput Comput Assist Interv. 2015, pp. 234–241. doi: 10.1007/978-3-319-24574-4_28.

[9] H. Chen et al. “DCAN: Deep Contour-Aware Networks for Object Instance Segmentation from Histology Images”. In: Med Image Anal 36 (2017), pp. 135–146. doi: 10.1016/j.media.2016.11.004.

[10] P. Naylor et al. “Segmentation of Nuclei in Histopathology Images by Deep Regression of the Distance Map”. In: IEEE Trans Med Imaging 38.2 (2019), pp. 448–459. doi: 10.1109/TMI.2018.2865709.

[11] J. W. Johnson. “Automatic Nucleus Segmentation with Mask-RCNN”. In: Advances in Computer Vision. 2020, pp. 399–407. doi: 10.1007/978-3-030-17798-0_32.

[12] U. Schmidt et al. “Cell Detection with Star-Convex Polygons”. In: Med Image Comput Comput Assist Interv. 2018, pp. 265–273. doi: 10.1007/978-3-030-00934-2_30.

[13] V. Kulikov and V. Lempitsky. “Instance Segmentation of Biological Images Using Harmonic Embeddings”. In: Proc IEEE Comput Soc Conf Comput Vis Pattern Recognit. 2020. 1904.05257.

[14] F. A. Guerrero Peña et al. “J Regularization Improves Imbalanced Multiclass Segmentation”. In: Proc IEEE Int Symp Biomed Imaging. 2020, pp. 1–5. doi: 10.1109/ISBI45749.2020.9098550.

[15] K. Löffler, T. Scherr, and R. Mikut. A Graph-Based Cell Tracking Algorithm with Few Manually Tunable Parameters and Automated Segmentation Error Correction. bioRxiv. 2021. doi: 10.1101/2021.03.16.435631.

[16] M. Maška et al. “A Benchmark for Comparison of Cell Tracking Algorithms”. In: Bioinformatics 30.11 (2014), pp. 1609–1617. doi: 10.1093/bioinformatics/btu080.

[17] V. Ulman et al. “An Objective Comparison of Cell-Tracking Algorithms”. In: Nat Methods 14 (2017), pp. 1141–1152. doi: 10.1038/nmeth.4473.

[18] D. Kingma and J. Ba. “Adam: A Method for Stochastic Optimization”. In: International Conference on Learning Representations. 2015. 1412.6980v9.

[19] L. N. Smith. “Cyclical Learning Rates for Training Neural Networks”. In: IEEE Winter Conf Appl Comput Vis. 2017, pp. 464–472. doi: 10.1109/WACV.2017.58.

[20] L. Liu et al. “On the Variance of the Adaptive Learning Rate and Beyond”. In: International Conference on Learning Representations. 2020. 1908.03265v3.

[21] M. Zhang et al. “Lookahead Optimizer: k steps forward, 1 step back”. In: Adv Neural Inf Process Syst. Vol. 32. 2019, pp. 9597–9608. url: https://proceedings.neurips.cc/paper/2019/file/90fd4f88f588ae64038134f1eeaa023f-Paper.pdf.

[22] H. Yong et al. “Gradient Centralization: A New Optimization Technique for Deep Neural Networks”. In: (2020). 2004.01461.

[23] I. Loshchilov and F. Hutter. “SGDR: Stochastic Gradient Descent with Warm Restarts”. In: International Conference on Learning Representations. 2017. 1608.03983.

[24] P. Matula et al. “Cell Tracking Accuracy Measurement Based on Comparison of Acyclic Oriented Graphs”. In: PLoS One 10.12 (2015), pp. 1–19. doi: 10.1371/journal.pone.0144959.

[25] K. Löffler and T. Scherr. KIT-Sch-GE(2). Cell Tracking Challenge algorithm description. 2021. url: https://public.celltrackingchallenge.net/participants/KIT-Sch-GE%20(2).pdf.

[26] R. Hataya et al. “Faster AutoAugment: Learning Augmentation Strategies Using Backpropagation”. In: Comput Vis ECCV. 2020, pp. 1–16. doi: 10.1007/978-3-030-58595-2_1.

